# Mesoscopic patterns of functional connectivity alterations in autism by contrast subgraphs

**DOI:** 10.1101/2022.11.19.517174

**Authors:** Tommaso Lanciano, Giovanni Petri, Tommaso Gili, Francesco Bonchi

## Abstract

Despite the breakthrough achievements in understanding structural and functional connectivity alterations that underlie autism spectrum disorder (ASD), the exact nature and type of such alterations are not yet clear due to conflicting reports of hyper-connectivity, hypo-connectivity, and –in some cases– combinations of both. In this work, we approach the debate about hyper- vs hypoconnectivity in ASD using a novel network comparison technique designed to capture mesoscopic-scale differential structures. In particular, we build on recent algorithmic advances in the sparsification of functional connectivity matrices, in the extraction of contrast subgraphs, and in the computation of statistically significant maximal frequent itemsets, and develop a method to identify mesoscale structural subgraphs that are maximally dense and different in terms of connectivity levels between the different sets of networks.

We apply our method to analyse brain networks of typically developed individuals and ASD patients across different developmental phases and find a set of altered cortical-subcortical circuits between healthy subjects and patients affected by ASD. Specifically, our analysis highlights in ASD patients a significantly larger number of functional connections among regions of the occipital cortex and between the left precuneus and the superior parietal gyrus. At the same time, reduced connectivity characterised the superior frontal gyrus and the temporal lobe regions. More importantly, we can simultaneously detect regions of the brain that show hyper and hypo-connectivity in ASD in children and adolescents, recapitulating within a single framework multiple previous separate observations.

## 1. Introduction

Autism spectrum disorder (ASD) is associated with disrupted or altered brain connectivity (Kana et al., 2014b) at both structural (Vissers et al., 2012) and functional (Hull et al., 2017a) levels. Deviations of the functional connections among brain regions from the normal pattern of connectivity are associated with several functional impairments (Abrams et al., 2013; Supekar et al., 2013). Specifically, alterations of the functional connectivity strength among brain regions support a variety of social (Gotts et al., 2012; Abbott et al., 2016), cognitive (Nair et al., 2015; Farrant and Uddin, 2016), and sensorimotor (Rudie et al., 2013; Di Martino et al., 2014) functions in ASD patients. Indeed, experimental evidence exists of a correlation between functional connectivity perturbations and ASD symptom severity (Fishman et al., 2014) and adaptive behaviour (Plitt et al.,2015). Despite the incredible results achieved in understanding structural and functional connectivity alterations that underlie core and associated features of ASD, the exact nature and type of such alterations are not completely clear due to conflicting reports of hyper-connectivity (Chien et al., 2015;Ray et al., 2014), hypo-connectivity (Just et al., 2012a), and –in some cases– combinations of both (Coben et al., 2014).

The techniques used to investigate the pathological functional connectedness of the brain typically deal with correlation networks built from the functional time series extracted from brain regions, whose spatial resolution can vary on a large mesoscopic scale (Khan et al., 2013). The results of network analyses at different scales can find similar topological properties across scales (van den Heuvel et al., 2015). The energetic and spatial constraints that shape network structure at the scale of brain regions and areas work similarly at the cellular level (Henriksen et al., 2016). Nonetheless, the function of network nodes and node clusters likely depends critically upon the scale at which a network is constructed and analyzed. Accordingly, we might expect networks to be optimized to perform scale-specific functions (Yuan et al., 2022), and focusing on a particular scale gives a unique insight into the network architecture underpinning those functions. Therefore, the open problem is finding a coherent approach to highlight connectivity alterations across multiple scales while retaining a global perspective on the network level.

Against this background, identifying specific neuroimaging-based biomarkers for ASD, especially ones that could be related to symptom severity, is still a challenging task, usually requiring large-scale datasets and analyses (Abraham et al., 2017;Doyle-Thomas et al., 2015). The reason is that, in addition to the issues described above, to be effective such biomarkers have not only to reach statistical significance but also to be stable across multiple datasets, experimental designs and subjects (Mueller et al., 2013). Similarly to other alterations of functional connectivity (Gheiratmand et al., 2017), existing methods rely either on the identification of network features specific to the ASD spectrum (Yerys et al., 2017) or on opaque neural network techniques (Sherkatghanad et al., 2019; Aghdam et al.,2018).

In this work, we approach the debate about hyper- vs hypoconnectivity in ASD, taking on a mesoscopic approach based on recent advances in network comparison techniques. We build on top of the recent proposal by (Lanciano et al., 2020) to detect contrastive subgraphs in terms of density between two groups of graphs obtained from resting state fMRI data. The method outputs group-level *contrast subgraphs*, which are maximally different, in terms of connectivity level, between the brain networks of typically developed (TD) individuals and ASD patients. We equip our method with simultaneous mining of different subgraphs exhibiting this property, making it able to provide a variety of findings at the same time. For example, we show here that, under these goggles, it is possible to reframe previous results on network architectures in ASD subjects in terms of a complex interplay between hyper- and hypo-connectivity, which evolves across age and relates deeply to individual differences. In addition to the group-level discrimination, the information obtained from contrast subgraphs can be used to build individual-level subgraphs, which can then be studied and related to individual phenotypical properties, for example, cognitive and social performances.

## 2. Methods

### 2.1. Subjects

Resting-state fMRI data from 57 males with ASD (15 children, 42 adolescents) and 80 typically developed (TD) (17 children, 63 adolescents) males were acquired from the Preprocessed Connectomes Project (Craddock et al., 2013). The data had been obtained from ABIDE (Martino et al., 2013), and preprocessed using the Data Processing Assistant for Resting-State fMRI (DPARSF) (Yan, 2010). Participants were excluded if mean framewise displacement (FD) during the resting-state fMRI scan was greater than 0.10 mm, and the percentage of data points exceeding 0.10 mm was greater than 5%. Groups were matched for age, IQ, mean FD, and the percentage of data points exceeding 0.10 mm. ASD diagnoses were confirmed using ADOS (Lord et al., 2000) and/or the Autism Diagnostic Interview-Revised (ADI-R; (Lord et al., 1994)). Participant characteristics are described in Appendix A.1. Written informed consent or assent was obtained for all participants in accordance with respective institutional review boards.

### 2.2. fMRI acquisition and preprocessing

Information about scanner types and parameters can be found on the ABIDE website.^1^ We carried out the following DPARSF preprocessing steps: slice timing correction, motion correction, realignment using a six-parameter (rigid body) linear transformation with a two-pass procedure (registered to the first image and then registered to the mean of the images after the first realignment). Individual structural images (T1-weighted MPRAGE) were co-registered to the mean functional image after realignment using a 6 degrees-of-freedom linear transformation without re-sampling. The transformed structural images were segmented into grey matter (GM), white matter (WM) and cerebrospinal fluid (CSF) (Ashburner and Friston, 2005) and nuisance parameters were regressed out (including 24 motion parameters, WM and CSF signals, linear and quadratic trends, and the global signal) (Satterthwaite et al., 2013). Temporal filtering (0.01 - 0.1 Hz) was performed on the time series.

The Diffeomorphic Anatomical Registration Through Exponentiated Lie algebra (DARTEL) tool (Ashburner, 2007) was used to compute transformations from individual native space to MNI space. We chose to use data that had the global signal regressed out, as this step has been shown to help mitigate differences across multiple sites (Power et al., 2014). Furthermore, it has been shown recently that global signal regression attenuates artifactual changes in BOLD signal that are introduced by head motion (Ciric et al., 2017; Power et al., 2017). The time series of 116 regions of interest (ROIs) from the Automated Anatomical Labeling (AAL) atlas (Tzourio-Mazoyer et al., 2002) were obtained. Additional details of the fMRI preprocessing steps can be found on the Preprocessed Connectomes Project website ^2^.

For any participant, we computed standard functional connectivity matrices from the preprocessed timeseries using Pearson’s correlation coefficient (Bassett and Sporns, 2017) (Figure 2A). We then sparsified each matrix using recent network-theoretical methods (SCOLA (Masuda et al., 2018; Kojaku and Masuda, 2019)), to obtain the individual sparse weighted network (with densities typically *ρ* < 0.1 consistent with standard sparsification methods (Hermundstad et al., 2013)).

**Figure 1:**
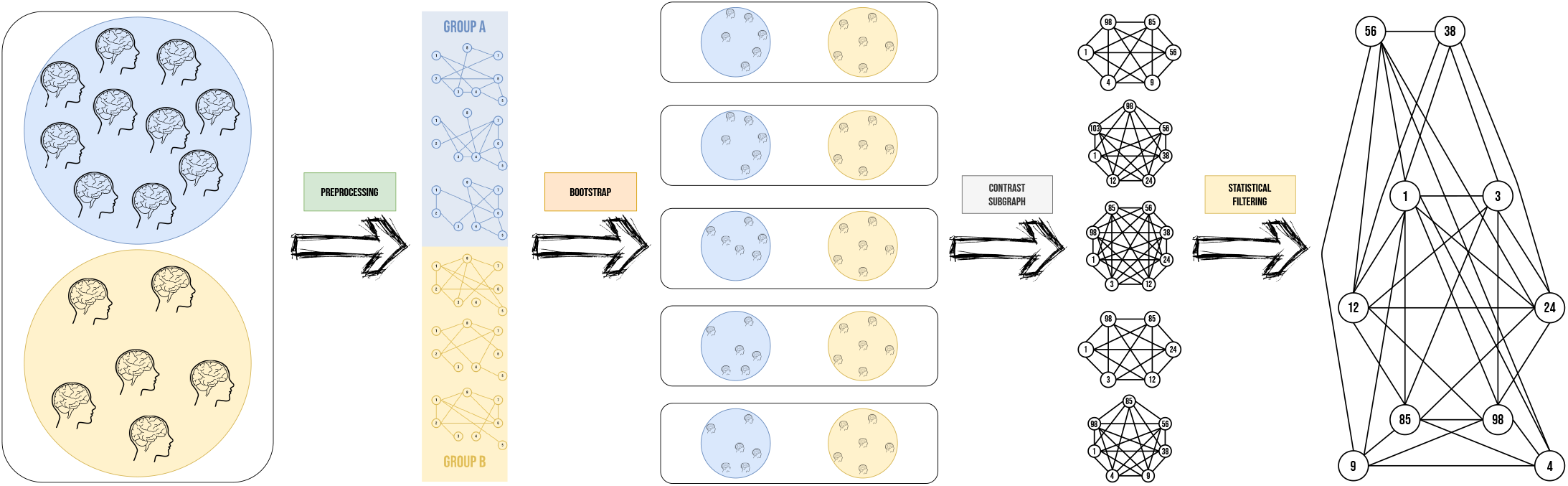
Our proposed pipeline. After an initial preprocessing of the fMRI scans that leads to the generation of the brain networks, we implement a bootstrap scheme that computes multiple contrast subgraphs, which are finally filtered to retain only those statistically significant. Each step of the pipeline is depicted, in finer details, in Figure 2.

**Figure 2:**
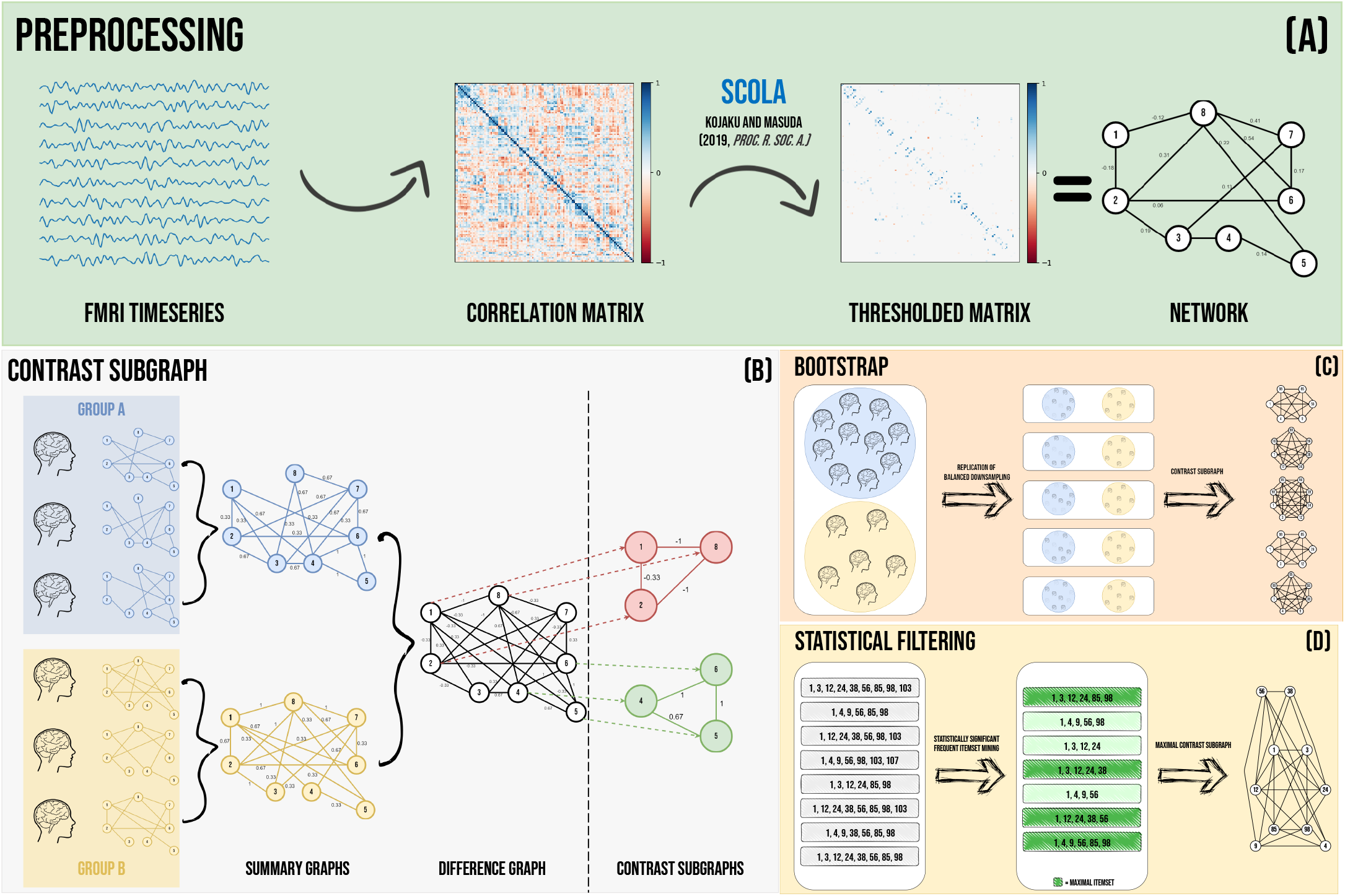
Details of our proposed pipeline. (A) fMRI preprocessing: We use BOLD timeseries to construct functional connectivity (FC) matrices, which we then sparsify using state-of-art network filtering techniques (Masuda et al., 2018; Kojaku and Masuda, 2019) to obtain sparse functional networks. (B) Computation of the contrast subgraph (Lanciano et al., 2020): Given the functional networks for two subject groups, the first step is to reduce both of them to their relative *summary graph*. Once both groups have been reduced, an optimisation problem is defined over the *difference graph*. Finally, we can solve and refine the optimisation problem with a local search approach. The final result is a set of nodes, dubbed *contrast subgraph*. In this instance, the two solutions make it clearer the approach followed: the set of nodes {1, 2, 8} is sparsely connected in the Group A, while it shows a large number of connections in the graphs of Group B. The set of vertices {4, 5, 6} shows the opposite pattern: density in the observations of Group *A* and sparsity in those of Group *B*. (C) Bootstrap of solutions: to compare unbalanced groups, we apply a down-sampling of the most populated group, such that it makes the groups balanced. For each resampled reduced dataset, we recompute the contrast subgraph as in (B), adopting as a value of *α* the one expressed by the proposal in (De Vico Fallani et al., 2017). (D) Alignment of bootstrap solutions: starting from the solutions obtained in the Bootstrap pipeline, we compute the statistically significant (overrepresented) maximal frequent itemsets (Kirsch et al., 2012), i.e. those that are not a subset of any other itemset. We compute the final solution as the union of these statistically significant maximal frequent itemsets (set of nodes) and the corresponding set of edges.

### 2.3. Contrast subgraphs extraction

We make use of a dataset 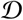 of functional networks, where the *i*-th observation corresponds to the sparse weighted network of the *i*-th individual, each one defined over the same set of nodes *V* (i.e., the 116 regions of the AAL atlas). 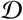 can be divided into two cohorts *c* = {children, adolescents}, each one composed of individuals belonging two one of the groups *g* = {TD, ASD}. Our proposal aims to detect multiple sets of regions of interest (ROIs) that, given a single cohort, simultaneously show hyper-connectivity in one group and hypo-connectivity in the other group. This can be achieved by appropriately generalizing the methodology proposed by (Lanciano et al., 2020) to fit our purposes. A summary of our proposed pipeline is depicted in Figure 1 while each step is represented in detail in Figure 2.

Given a cohort of individuals, for each group, we combine the group’s networks in a single graph, dubbed *summary graph:* this step is crucial because it compresses in a single observation the peculiarities contained across the group of networks. Then, considering the two summary graphs, we combine them in a *difference graph*, a new network whose edges have weights equal to the difference of the weights of the two summary graphs. Once the *difference graph* was computed, an optimization problem is defined on it, whose solution is the *contrast subgraph*, i.e. the set of ROIs that maximizes the difference in terms of density between the graphs of two groups. A general summary of the whole pipeline is depicted in Figure 2B. In Supplementary Material (Appendix A.2), we provide further details on the technical implementation of the optimization.

We note here that while the extraction of a contrast subgraph is defined at the group level, the relative individual subgraphs induced are different across individuals in the same group, which in turn provide subject-level information that can be exploited (as we will do, for example, in Section 3.4).

### 2.4. Computation and robustness of contrast subgraphs

Towards our objective, the pipeline described in the previous section has three main limitations that need to be taken into account:

1. the potential group imbalance in the dataset: imbalances between the number of subjects in each class can affect the optimization results, biasing it toward one of the classes;
2. the dependence of the contrast subgraph extraction on an accuracy parameter *α* (see Appendix A.2 for formal definition): indeed, depending on the value of *α*, it is possible to exclude potential ROIs that are contrastive but not the most contrastive one;
3. the presence of multiple subgraphs exhibiting the contrast property, that are not detected by the optimization problem proposed in (Lanciano et al., 2020).

Therefore, starting from the algorithm provided in (Lanciano et al., 2020), we employ an enhance to the existing pipeline to efficiently address all these limitations. To address the first issue we employ a subsampling scheme to correct class imbalances. In particular, for a dataset 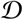 with unbalanced groups 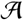 and 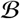, we perform a bootstrap procedure (Figure 2C), by subsampling a number 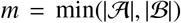 of subjects from each group, and repeating the contrast subgraph extraction for each subsample. Thanks to this procedure, we are also able to tackle the third issue, producing a larger number of solutions. To address the second issue, we relate our choice of the value of the parameter *α* to follow the proposal of (De Vico Fallani et al., 2017), which provides a topological criterion to threshold a graph in such way there is balance in the trade-off between efficiency of a network and its wiring cost. Therefore, we set *α* equal to this value, so that the positive contribution to the objective function is brought only from edges whose weight is greater than the threshold.

Finally, to solve the third issue, we rely on the bootstrap scheme proposed above, and test the general representativeness of the obtained solutions by reconciliating the multiple subgraphs using a filter based on the Frequent Itemset Mining paradigm (Figure 2D). The problem can be formalized as follows: we are given a set of transactions (contrast subgraphs), each one containing a set of different items (nodes), and we want to detect those items who are often co-occurring (those groups of ROIs that are often contrastive between the 2 different classes). Therefore, considering the different solutions computed, we mine the most frequent contrast subgraphs that are also statistically significant (such that False Discovery Rate ≤0.01), exploiting the proposal of (Kirsch et al., 2012), and retain only those that are maximals (i.e., sets that are not the subset of any other set in the collection). In this way, we are able to address the last limitation previously described, and to obtain a single solution by considering the union of these statistically significant maximal frequent itemsets (set of nodes), and the corresponding set of edges. Further technical details of these procedures are provided in Appendix A.3.

## 3. Results

### 3.1. Contrast subgraphs classify subject group

Following the methods described above, we extracted the two edge-sets that define the contrast subgraphs encoding the main network-level differences between the two groups. While these edge-sets are determined at the group level, it is naturally possible to observe fluctuations in the individual-induced subgraphs –the graph obtained by projecting the subgraph of the individual functional connectivity on the edges induced by such dense sets of nodes. This leads to variable levels of hyper/hypo-connectivity, measured in terms of the weights’ sum of edges in the induced subgraph across subjects. Indeed, by considering these overall connectivity levels (as described in Methods and Fig. 2), we can obtain a separation boundary to separate subjects in terms of their induced graph density that provides a simple rule to classify subjects in one of two groups. While identifying this boundary is not the primary aim of the contrast subgraph extraction method, the differences in densities between groups (Fig. 3A and 3B for children and adolescents respectively) are clear. To confirm this and validate the goodness of our solution, we solve the classification task described above over any repetition of the bootstrap, running a linear SVM over the whole dataset to find a linear separation bound between the two groups, which can classify with large accuracy. Interestingly, the separation based on the contrast subgraphs is more effective in children (accuracy 0.80±0.06) than adolescents (accuracy 0.68 ±0.04).

**Figure 3:**
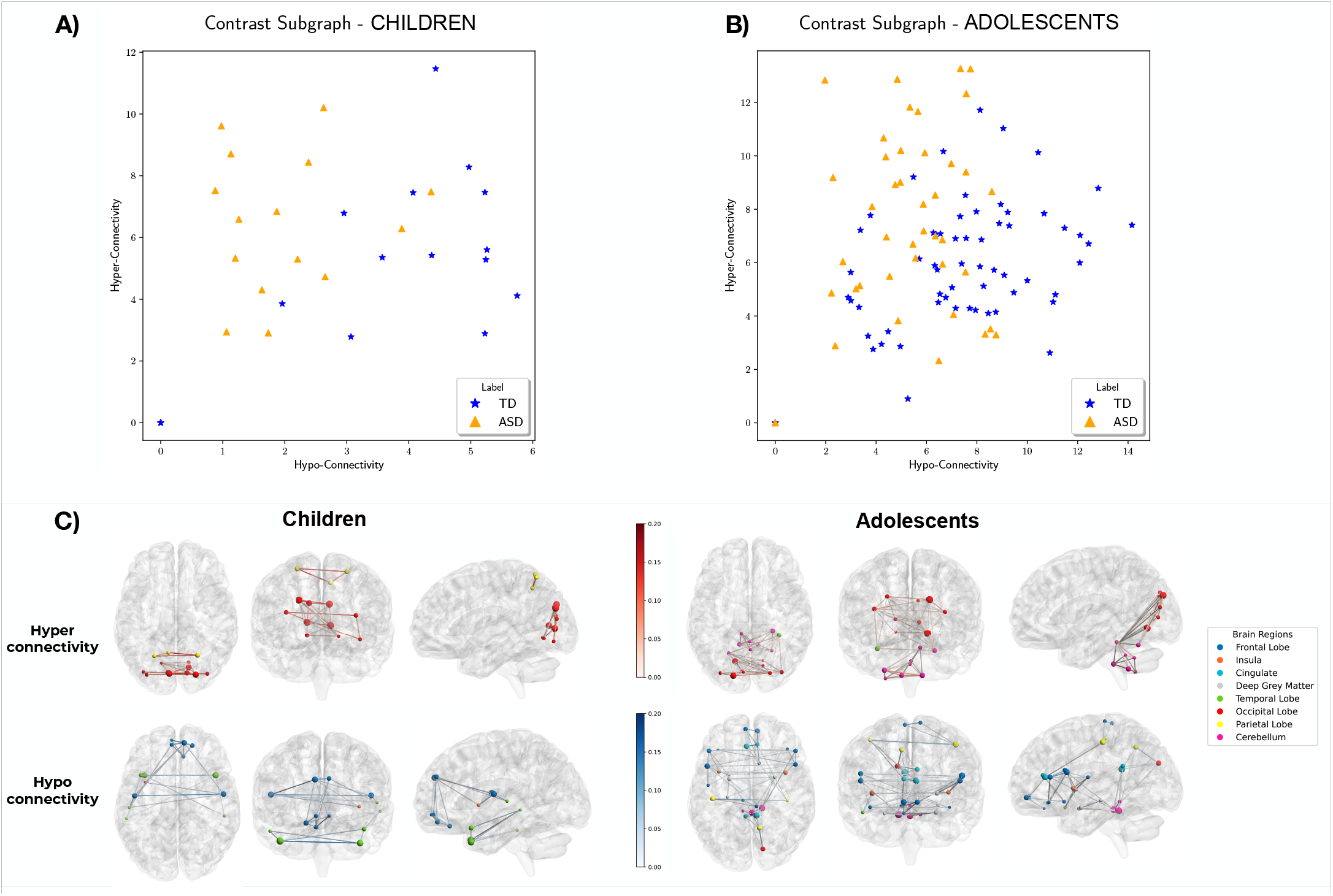
(A) Level of hyper-connectivity and hypo-connectivity for Children (left) and Adolescents (right). Connectivity is expressed as the number of edges inside the contrast subgraph. We refer to hyper-connectivity for the solution denser for ASD and sparser in TD (and vice versa for hypo-connectivity). See main text for accuracy reports. (B) Visualization of the contrast subgraphs for the two age groups, children (left) and adolescents (right). Red (blue) edges represent edges present in the contrast subgraph for the ASD subjects (TD subjects).

### 3.2. Node-level description of contrast subgraphs

Below we report in the extended form our results for hyper- and hypo-connectivity patterns in ASD individuals (as compared to TD ones), separately for children and adolescents. All the reported regions and substructures have been statistically validated for the contrast generated by the two classes, according to the U-Test (*p* < 0.05). If not otherwise indicated, brain regions reported must be considered as bilaterally involved. Figure 3C depicts the contrast subgraphs for the two age groups (left, children; right, adolescents), highlighting the edges retained in the Contrast Subgraphs that are hyper-(hypo-)expressed in ASD subjects as compared to TD ones (3C top and bottom row respectively).

#### Children

##### Hypo-connectivity

We find the subgraph mostly localized inside the Frontal Lobe. In particular, we find a central role of the Superior Frontal Gyrus (Medial), that shows, bilaterally, hypo-connectivity with two different sub-modules: (i) Orbital part of the Superior Frontal Gyrus, Medial Orbital part of the Superior Frontal Gyrus and Right Rectus; (ii) Rolandic Operculus, Left Insula and Left Superior Temporal Gyrus. Inside the Temporal Lobe, in particular between the Temporal Pole of the Superior and Middle Temporal Gyrus, and their connections with Right Inferior Temporal Gyrus and Left Middle Temporal Gyrus.

#### Children

##### Hyper-connectivity

We find localized structures: between Middle Occipital Gyrus and Left Inferior Occipital Gyrus; between Calcarine, Cuneus, Lingual and Right Superior Occipital Gyrus; and between the Superior Parietal Gyrus and the Left Precuneus.

#### Adolescents

##### Hypo-connectivity

We find, in this case, a huge contrastive structure involving di*ff*erent patterns of disconnection that include: (i) Inferior Frontal Gyrus (Triangular, Opercular and Orbital parts), the Insula, Left Putamen, Left Rolandic Operculus and the Temporal Pole of the Right Temporal Gyrus; (ii) Amygdala and Left Hyppocampus; (iii) Posterior Cingulate Gyrus, Right Cuneus and Right Precuneus; (iv) Vermis 3 and Left Cerebellum 3. Each submodule shows a significant disconnection from the cerebellar Vermis 1 and 2 and the Right Cerebellum 3. We also find a unique contrast subgraph composed of: the Postcentral and the Anterior Cingulate Gyrus, the Orbital Part of the Middle Frontal Gyrus, the Right Olfactory cortex, the Paracentral Lobule and the Orbital Part of the Middle Frontal Gyrus.

#### Adolescents

##### Hyper-connectivity

The Contrast subgraphs are localized in the Occipital Lobe (Superior and Middle Occipital Gyrus, Lingual, Calcarine and Left Cuneus) with connections with the Right Fusiform and the Left Cerebellum 6. We also find nodes within and between Cerebellum (Left 3, Right 8, 9, Right 10) and Vermis (9 and 10).

### 3.3. Mesoscopic differences in lobe integration across age-groups

To summarise the information and to understand how the large-scale integration patterns differ across conditions, we coarse-grain the individual induced subgraphs from nodes to brain lobes (see Appendix A.4). We then select the edges expressed in a significantly stronger way in ASD than in TD subjects for each age group (see Appendix A.4), which we denote as *hyper-expressed*, and those that are instead weaker in adolescents versus children, which we denote as *hypo-expressed*.

In Figure 4A we show the adjacency matrices of the lobe-level induced graphs, divided by hyper- and hypo-expression for the four conditions. The first observation is that, at the mesoscopic level, both hyper- and hypo-expressed ASD-induced graphs show significant connectivity between occipital and cerebellar regions, which is instead absent in the case of TD subjects. More in detail, we also see that the amount of hypo- and hyper-expression is larger for ASD than for TD graphs between the occipital and temporal lobes and within the cerebellum. Together these observations suggest a nuanced reorganization of the ASD subjects’ brains from childhood to adolescence than in TD subjects. In particular, we do not only observe more or less connectivity but also find a combination of both effects simultaneously. We also find that lobes differ in the evolution of their self-interaction across age and condition (Fig. 4B). In particular, for the cingulate lobe, deep grey Matter, and insula, we find a significant increase (linear regression, *p* < 0.05 on all coefficients) of lobe integration with age for TD subjects, which is instead absent in ASD individuals. The parietal and temporal lobes show conflicting trends: in the former, self-integration significantly increases for TD subjects and decreases for ASD subjects, while for the latter, the trends are reversed. Finally, we find that the cerebellum is progressively more integrated with age in both groups, but the increase of integration with age is significantly larger for ASD subjects. For the frontal and occipital lobes, we did not find any significant group-level effect with age for either ASD or TD subjects. Overall, our analysis shows that, whereas TD subjects develop greater local integration within lobes as they grow, ASD subjects maintain a more distributed architecture with the notable exceptions of the cerebellum and the temporal lobe.

**Figure 4:**
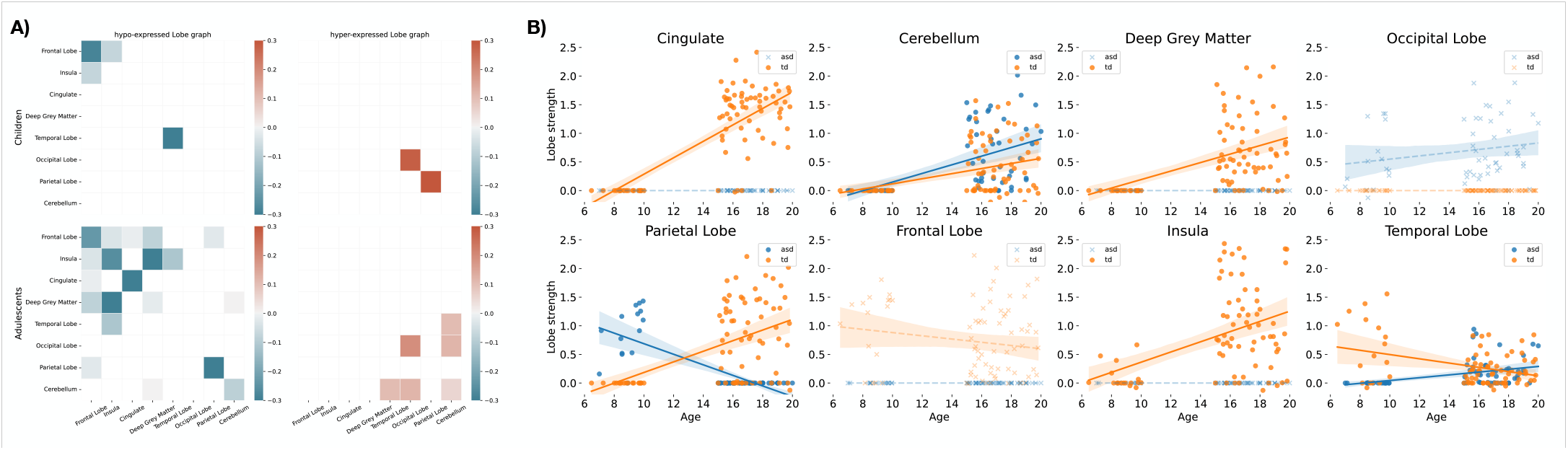
(A) Hyper and hypo-integration of lobes in ASD subjects vs TD subjects for the two age groups. We select either hypo- or hyper-expressed edges in the induced subgraphs corresponding to the various conditions and then coarse-grain the resulting networks to the lobe level to highlight the mesoscopic differences in the integration patterns across groups. Positive/negative values imply stronger edges within/between lobes (by coarse graining over the corresponding regions) in adolescent/children subjects. Overall, we find stronger interactions between occipital and temporal lobes in ASD for TD, larger differences within the cerebellar areas, and –finally– between the cerebellum and the occipital lobe. (B) Age dependence of lobe self-strength in induced lobe graphs. Full dots represent quantities for which the linear regression coefficients are significant (*p* ≤ 0.05), while crosses show results for quantities for which we find no significant trend. For most lobes, we find a significant increase with the age of lobe-level self-strength for TS subjects. In contrast, ASD subjects only show significant increases in the temporal lobe and cerebellum..

### 3.4. Induced contrast subgraphs correlate with individual phenotypes

The contrast subgraphs discussed so far describe the salient features of each subject group and condition. However, inspired by the previous results, we wonder whether any differences observed at the lobe-level integration correlate with individual performance. We can investigate this by considering changes in individual architectures within the same group, for example, whether these sets can highlight individual phenotypical differences. To do this, we consider the induced contrast subgraphs at the individual level: given a cohort of patients, once computed the contrast subgraph *C*^*g*_1_,*g*_2_^, i.e., the set of nodes whose induced subgraph is denser in group *g*_1_ and sparser in group *g*_2_, we consider, for any individual *i* of the cohort, the subgraph 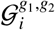 obtained by restricting the functional connectivity network of *i* to the edge-set induced by the node-set *C*^*g*_1_,*g*_2_^.

We then investigate whether simple graph metrics, i.e. lobe integration strength and lobe-lobe integration strength, correlate with the subjects’ phenotypes as measured by standard performance scales (Lord et al., 1994, 2000). In Figure 5 we report the statistically significant (*p* < 0.05 before Bonferroni correction) relationships we obtained. We find that cerebellar and occipital lobe integrations are negatively correlated with standard intelligence scores (F/VIQ), and that the strength of the interaction between the occipital and temporal lobes in the lobe-level graphs associates with more severe social symptoms.

**Figure 5:**
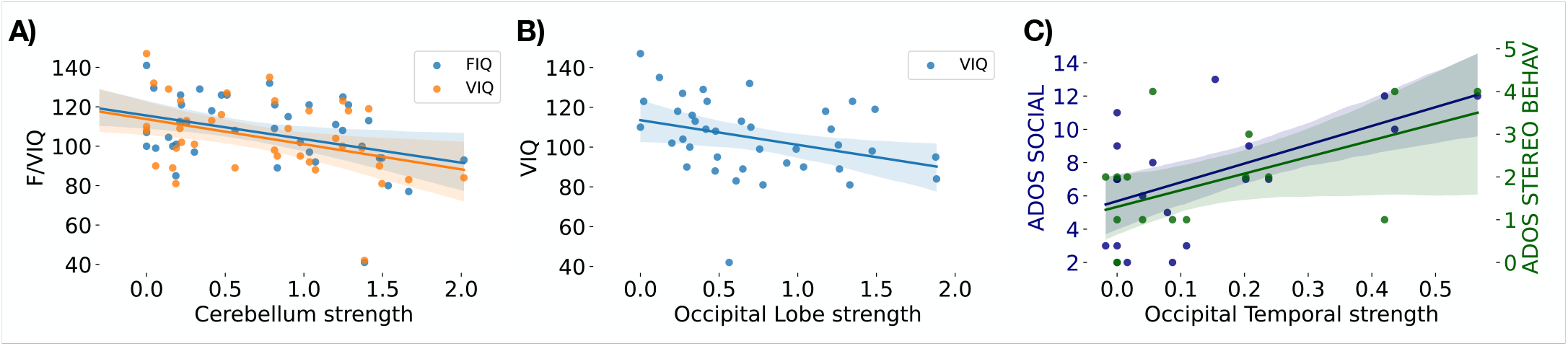
Association between lobe strengths and lobe interactions with phenotypes in ASD adolescents. We report i) the significant (*p* < 0.01) associations between cerebellar (A)) and occipital (B) lobe strength, respectively, with intelligence scores, FIQ and VIQ; and ii) the significant association of the interaction strength between occipital and temporal lobes (C) with social scores (ADOS Social and ADOS Stereo Behaviour)(Craddock et al., 2013).

## 4. Discussion

Using contrast subgraph analysis, we elucidated atypical resting-state brain functional connectivity underlying the symptoms of ASD both in children and adolescents. Specifically, we analyzed a large rs-fMRI dataset from the ABIDE database to determine whether neurophysiological changes typical of the disease are associated with cortico-cortical and cortico-subcortical dysfunctional connectivity in the brains of males with ASD. Our findings indicated that children of the ASD group exhibited a significantly larger number of functional connections among regions of the occipital cortex and between the left precuneus and the superior parietal gyrus. At the same time, reduced connectivity characterized the superior frontal gyrus as well as regions of the temporal lobe. Conversely, adolescents of the ASD group showed hypo connectivity of Vermis 1, Vermis 2 and right III lobule of the cerebellum, while hyper-connectivity was found between regions of the occipital cortex and the left IV lobule of the cerebellum and the right fusiform, as well as among the left III, the right 8, 9 and 10 lobules of the cerebellum and Vermis 9 and 10.

The method proposed here captured both hyper-connectivity and hypo-connectivity across the whole brain, as opposed to common functional connectivity approaches that typically are only able to unveil only one direction of alteration or a subset of brain regions involved (Kana et al., 2014a; Hull et al., 2017b;King et al., 2019). This paper offers strong evidence that autism is characterized by atypical large-scale brain functional connectivity in both directions (Uhlhaas and Singer, 2012). Previous human and model neuroimaging studies reported a variety of functional architectures specific to ASD (Just et al., 2012b; Anderson et al., 2012; Yahata et al., 2016). Most of them did not directly allow the simultaneous observation of under- and over-expressed connectivity of regions across the whole brain in ASD individuals, both at the regional and lobe level and of their evolution during development.

On the one hand, our results show that along with the development of the functional connectivity over-expression pattern in ASD patients include regions of the occipital cortex both in children and in adolescents, as already observed in previous rs-fMRI studies (Keown et al., 2013; Nair et al., 2018). According to the weak central coherence model (Frith, 2003; Happé and Frith, 2006), overstimulated and underselective visual processing areas may dominate high-order cognitive processes in autism. This can lead to the reduced ability in contextual information integration during the execution of complex perceptual and executive tasks (Belmonte et al., 2004) and can be related to the increased connectivity of visual cortices (Groen et al., 2010; Shen et al., 2012; Jao Keehn et al.,2017). We also find increased connectivity between visual regions across hemispheres, suggesting enhanced perceptual processing, as observed in (Barbeau et al., 2015; Clawson et al.,2015; Jao Keehn et al., 2019) together with the intact interhemispheric transfer of visual information.

On the other hand, we find differential hyper-connectivity in children and adolescent patients. Specifically, while the increased bilateral connectivity in children characterized regions belonging to the parietal cortex, adolescent ASD patients were characterized by increased integration of the cerebellum. Hyper-connectivity found between superior parietal gyrus and precuneus might reflect altered patterns of signal fluctuations (Uddin et al., 2013) in the interaction between the networks these regions originate from (fronto-parietal and default mode network), that, in turn, may impair cognitive processing. Increased coordination with nonessential regions may introduce low-level cross-talk and spoiling signal across primary network components (Belmonte et al., 2004). These results, therefore, constitute a possible source of the widely observed decreased connectivity within the Default Mode Network (Lynch et al., 2013; Washington et al., 2014; Padmanabhan et al., 2017;Kotila et al., 2021), which may stem from the disturbing abnormal connectivity that one of its hubs (precuneus) has with different and unrelated regions (bilateral superior frontal gyrus).

Remarkably, atypical visual exploration of both social and nonsocial scenes is often reported in ASD patients with less precise and longer saccades, potentially reflecting difficulties in eyes movement control (Kovarski et al., 2019), exerted by the cerebellar lobules VIII-X and uvula/nodulus (Vermis IXX) (Vahedi et al., 1995; Stoodley and Schmahmann, 2010). The increased connectivity of the cerebellar lobules VIII-X and Vermis IX-X observed in our study might relate to the frequently reported reduction in gamma-aminobutyric acidergic (GABAergic) Purkinje cells in ASD (Bailey et al., 1998;Rubenstein and Merzenich, 2003). These cerebellar neurons send inhibitory projections to the deep cerebellar nuclei (the output nuclei of the cerebellum) and the posterior lobe of the Vermis. Loss of these neurons is thought to lead to disinhibition of the deep cerebellar nuclei and of the uvula/nodulus lobules (Cerliani et al., 2015; Belmonte et al., 2004), which could explain the observed increased cerebellar integration and consequent abnormal eyes movement control in ASD patients (Trimarco et al., 2021).

Patterns of regional hypo-connectivity in ASD have been widely reported (Moseley et al., 2015; Roy and Uddin, 2021;Cheng et al., 2015; Anderson et al., 2011; Nomi and Uddin,2015). Accordingly. At the same time, children showed reduced connectivity in frontal and temporal regions, and adolescent showed two heterogeneous patterns of hypo-connectivity: one involving cerebellar-subcortical, cuneo-cerebellar, fronto-cerebellar and fronto-parietal connectivity, the other involving anterior cingulate cortical connectivity with the parietal (postcentral gyrus) and frontal regions. The developmental trajectory of the illness across age is firstly characterized by the increased complexity of the disconnection patterns. In fact, children show a clear disgregation of the functional fronto-temporal network, already associated with impaired working memory processes (Urbain et al., 2016), language difficulties (Verly et al., 2014) and diminished preferential attention to social cues (Sperdin et al., 2018). In contrast, adolescents show functional network disconnections organized in a mosaic of modules.

Fronto-temporal (Alaerts et al., 2014; Malaia et al., 2020) and insular-limbic (Ebisch et al., 2011) dysconnectivity associated with emotion deficits, posterior cingulate-cuneus (Nair et al., 2020; Bathelt and Geurts, 2021) underconnectivity that reflects a slow and delayed maturation of the Default Mode Network associated with social impairments and cerebellar disconnections (Long et al., 2016; Van Overwalle et al., 2020) associated to a reduced social cognition, somatosensory and language skills have been separately reported in literature. Once the regions of interest were grouped in eight anatomical macroareas, we calculated, subject by subject, the average correlation within each one and tested the possible topology-phenotype covariance. We found that cerebellar and occipital cortices integration negatively correlated with standard measures of IQ (VIQ/FIQ) in ASD adolescents only, while no significant correlation was observed in children. Similarly, the increased connectivity of occipital and temporal areas was significantly associated with reduced social skills in ASD adolescents (ADOS Social, stereotypical behaviour scales). Besides the reduced sample size because of data homogeneity reasons, the absence of similar correlations in children may also suggest a complex role of developmental reorganization during the development of cognitive skills.

From a methodological perspective, it is important to highlight the mesoscopic nature of our approach. This is grounded in the optimization involved in extracting contrast subgraphs, which, while acting locally at the level of the single edge contribution (i.e. the difference in weight between the same edge in the two groups), obtains global solutions because the optimization takes into account the whole network. Conceptually, this is similar to what is done, for example, for network modularity (Brandes et al., 2008): in that case, however, the focus is on the modular density structure of a single network, not on the most discriminating subgraph between graphs representing two groups (or conditions). Crucially, and similarly to other mesoscopic observables, such as network modules (Sporns and Betzel, 2016; Esfahlani et al., 2021; Sigar et al., 2022) and holes (Petri et al., 2014; Ibáñez-Marcelo et al., 2019; Lee et al., 2012;Chung et al., 2019; Sizemore et al., 2018), the differences detected by contrast subgraphs pertain to the level of coordination among sets of nodes, rather than to the alteration of the integration patterns of specific nodes (Lin et al., 2014; Gallos et al.,2012).

Despite the method’s novelty and results described here, the contrast subgraph approach suffers from some limitations. First, because the method is intrinsically mesoscopic, it can detect localized differences between groups less. It should be adopted in situations where mesoscopic changes rather than specific regional ones are envisioned. Second, the method has a resolution parameter, *α*, that has a clear algorithmic interpretation, but that might be of difficult interpretation from a clinical point of view. However, combined with principled thresholding techniques, our bootstrapping approach can effectively buffer this problem. Third, in the optimization, we considered a simple difference between the summary graphs with a penalty term that does not depend on the individual edges nor the statistics of the graphs under comparison. This could be improved by substituting the penalty with one obtained from a network-generating model, e.g. a weighted configuration model Garlaschelli and Loffredo (2009); Voitalov et al. (2020).

## Data and code availability statement

The dataset analyzed in the present study, and the implementation of the algorithms are available by inquiry.

## Author contribution statement

**Tommaso Lanciano:** Methodology, Software, Investigation, Visualization, Writing-Original draft preparation. **Giovanni Petri:** Conceptualization, Methodology, Investigation, Visualization, Writing-Original draft preparation. **Tommaso Gili:** Methodology, Validation, Writing-Original draft preparation. **Francesco Bonchi:** Conceptualization, Methodology, Writing-Original draft preparation.

## Funding

The authors have no funding to disclose.

## Declaration of Competing Interest

The authors have no conflict of interest to disclose.

## Appendix A. Supplementary Material

### Appendix A.1. Participants

Table A.1 and Table A.2 report the information of the participants considered for all of the analysis in this work. For numerical variables, we report avg. value, standard deviation, min-max values and t-test result between the two groups. For categorical variables we report the absolute frequencies, and the chi-square test result between the two groups.

### Appendix A.2. Contrast subgraph extraction

We provide a generalized definition of the methodology employed in order to obtain the results reported in Section 3.

We consider a dataset 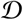 of observations, where the *i*-th observation corresponds to the pre-processed fMRI scan of the *i*-th individual. From any observation of 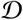 we are able to derive the relative brain network *G_i_* = (*V*, *E_i_*). The set of vertices *V* contains the brain regions, thus it is common to all the graphs, while the edge set *E_i_* represents connections between vertices in the observation graph *G_i_*. The dataset 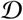 is divided in two groups: the *condition group* 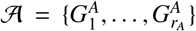 and the *control group* 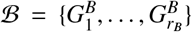. We aggregate the information in the groups 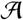 and 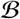 in two *summary graphs* 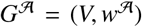 and 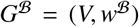, respectively, as follows. Given a group of networks 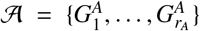, we define the *summary graph* 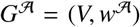 to be an undirected and weighted graphs, where 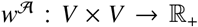 is a weight functions assigning a value to each pair of vertices. The role of 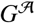 and 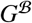 is to summarize in a single network the informations contained respectively in all the networks of 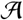 and 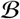. In this work, we employ the same weighting function of (Lanciano et al., 2020): given two

**Table A.1:** Participants information for the *children* cohort.

|  | Children |  |  |
| --- | --- | --- | --- |
|  | TD | ASD | Significance |
| **N** | 17 | 15 |  |
| **Age** | 8.767±0.935<br>[6.47-10] | 8.958±0.911<br>[7-9.95] | t = -0.564, p = 0.577 |
| **Gender** | 17 M | 15 M |  |
| **IQ** | 111.088± 9.681<br>[98-129] | 111.5± 23.574<br>[78.5-148] | t = -0.064, p = 0.949 |
| **Mean FD** | 0.053± 0.019<br>[0.025-0.094] | 0.052± 0.018<br>[0.026-0.083] | t = 0.123, p = 0.903 |
| **Perc FD** | 1.327± 1.321<br>[0-3.98] | 1.626± 1.325<br>[0-3.867] | t = -0.617, p = 0.542 |
| **Handedness** | 9 R<br>1 L<br>1 Ambi | 3 R<br>2 L<br>1 Ambi | chi = 2.039, p=0.361 |
| **Eyes Status** | 13 Open<br>4 Closed | 13 Open<br>2 Closed | chi = 0.08, p = 0.777 |
| **Site** | 1 MAX_MUN<br>3 STANFORD<br>6 NYU<br>3 KKI<br>1 UM.1<br>3 YALE | 0 MAX_MUN<br>1 STANFORD<br>9 NYU<br>1 KKI<br>3 UM.1<br>1 YALE | chi = 5.496, p = 0.358 |
| **Comorbidities** | 0 Yes<br>17 No | 8 Yes<br>7 No |  |
| **Currently Medicated** | 11 No<br>0 Yes | 9 No<br>5 Yes |  |
| **ADOS Total** | / | 11.182± 4.589<br>[7-21]<br>N=11 |  |
| **ADOS Comm** | / | 3.273± 1.863<br>[0-7]<br>N=11 |  |
| **ADOS Social** | / | 7.909± 3.26<br>[3-14]<br>N=11 |  |
| **ADOS Rrb** | / | 3.385± 1.862<br>[1-7]<br>N=13 |  |
| **ADI R Social** | / | 18.867± 5.188<br>[11-27]<br>N=15 |  |
| **ADI R Verbal** | / | 16.333± 5.055<br>[8-23]<br>N=15 |  |
| **ADI R Rrb** | / | 5.8± 2.4<br>[0-10]<br>N=15 |  |
| **SRS** | 14.111± 13.617<br>[2-50]<br>N=9 | 98.1± 21.366<br>[44-126]<br>N=10 | t = -9.546, p = 0 |

**Table A.2:** Participants information for the *adolescents* cohort.

|  | Adolescents |  |  |
| --- | --- | --- | --- |
|  | TD | ASD | Significance |
| **N** | 63 | 42 |  |
| **Age** | 17.128± 1.444<br>[15-19.83] | 17.062± 1.378<br>[15-20] | t = 0.232, p = 0.817 |
| **Gender** | 63 M | 42 M |  |
| **IQ** | 110.92± 11.962<br>[81-129] | 108.963± 18.005<br>[41-141] | t = 1.285, p = 0.202 |
| **Mean FD** | 0.048± 0.018<br>[0.024-0.094] | 0.055± 0.021<br>[0.018-0.098] | t = -1.599, p = 0.113 |
| **Perc FD** | 1.036± 1.287<br>[0-4.972] | 1.381± 1.378<br>[0-4.636] | t = -1.295, p = 0.198 |
| **Handedness** | 40 R<br>6 L<br>1 Ambi | 27 R<br>1 L<br>1 Ambi | chi = 1.939, p=0.379 |
| **Eyes Status** | 44 Open<br>19 Closed | 34 Open<br>8 Closed | chi = 1.099, p = 0.294 |
| **Site** | 2 PITT<br>8 UM.2<br>2 YALE<br>2 USM<br>5 SDSU<br>3 CALTECH<br>7 TRINITY<br>0 LEUVEN.1<br>14 NYU<br>5 LEUVEN.2<br>2 OLIN<br>13 UM.1 | 0 PITT<br>3 UM.2<br>2 YALE<br>4 USM<br>5 SDSU<br>1 CALTECH<br>6 TRINITY<br>5 LEUVEN.1<br>7 NYU<br>0 LEUVEN.2<br>3 OLIN<br>6 UM.1 | chi = 17.634, p = 0.09 |
| **Comorbidities** | 0 Yes<br>63 No | 4 Yes<br>38 No |  |
| **Currently Medicated** | 48 No<br>1 Yes | 22 No<br>13 Yes |  |
| **ADOS Total** | 2± 1<br>[1-3]<br>N=2 | 11.038± 3.684<br>[4-19]<br>N=26 |  |
| **ADOS Comm** | 0.5± 0.5<br>[0-1]<br>N=2 | 4± 1.643<br>[0-7]<br>N=20 |  |
| **ADOS Social** | 1.5± 0.5<br>[1-2]<br>N=2 | 7.15± 3.336<br>[2-13]<br>N=20 |  |
| **ADOS Rrb** | 0<br>[0]<br>N=1 | 3± 1.592<br>[0-6]<br>N=15 |  |
| **ADI R Social** | / | 19± 4.564<br>[11-28]<br>N=29 |  |
| **ADI R Verbal** | / | 15.034± 4.537<br>[2-23]<br>N=29 |  |
| **ADI R Rrb** | / | 5.621± 2.311<br>[2-12]<br>N=29 |  |
| **SRS** | 18.727± 11.993<br>[4-46]<br>N=11 | 87.857± 24.222<br>[42-123]<br>N=7 | t = -7.583, p = 0 |

vertices *u* and *v* in *V*, we define 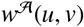 to be the *fraction* of graphs 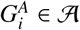 in which *u* is incident to *v*, that is,

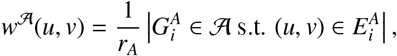

and similarly for 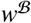. Note that according to this weighting function 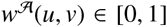 with 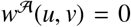 denoting the case in which there is no relationship (i.e., no edge) between *u* and *v* in 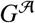. Given a subset of vertices *S* ⊆ *V*, we define

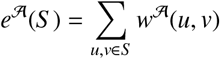

the sum of edge weights in the subgraph of 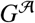 induced by the vertex set *S*, i.e. the graph that is made by the vertices in *S*, and those edges that connect any pair of vertices (*u*, *v*) ∈ *S*. Analogous definitions apply to 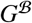.

The problem of extracting the contrast subgraph requires to find a subset of vertices whose induced subgraph is dense (i.e., has an high number of edges) in a summary graph 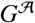 and sparse (i.e., has a low number of edges) in summary graph 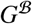.

#### Problem 1 (Contrast subgraph extraction)

*Given two sets of observation graphs, i.e., the condition group 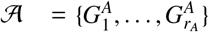 and the control group 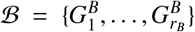, and corresponding summary graphs 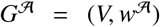 and 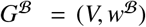, we seek to find a subset of vertices S*\* ⊆ *V that maximizes the* contrast-subgraph objective

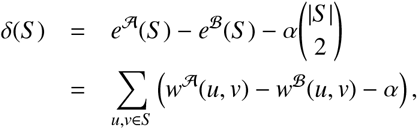

where 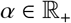 is a user-defined parameter

The objective function *δ*(*S*) is composed by two parts: 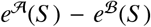 represents the contrast, i.e., the difference in terms of absolute number of edges inside the set of nodes between the 2 groups, while 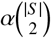, is a regularization term penalizing solutions of large size: in fact, given that a graph of size |*S*| can contain at most 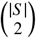 edges, it assign to any single edge included in the final solution a constant penalty α. In such a way, the magnitude of contrast has to be stronger than the penalty factor. Note that, to avoid the naïve solution, 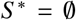, we have to ensure that 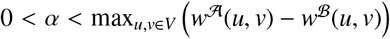, otherwise we would encounter the case in which every pair of vertices is detrimental for the objective function.

To solve Problem 1 is possible to exploit an algorithmic result by (Cadena et al., 2016) devised to solve the following problem.

#### Problem 2 (Generalized optimal quasi-clique)

*Given a graph G* = (*V*, *E*), *and functions w*(*u*, *v*) *and α*(*u*, *v*), *for each pair of vertices u*, *v* ∈ *V, find a subset of vertices S* ⊆ *V that maximizes*

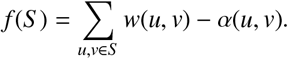

It is straightforward to see that, by setting *α*(*u*, *v*) = *α* and 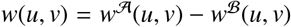 for each pair of vertices *u*, *v* ∈ *V*, Problem 1 can be mapped to Problem 2. Therefore, solving Problem 1 with 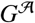 and 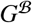 in input is equivalent to solve Problem 2 with 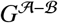 in input. We call 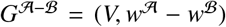 the *difference graph*.

Cadena et al. prove that Problem 2 is **NP**-complete and **NP**-hard to approximate within a factor 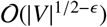 (Cadena et al.,2016, Theorem 1). Then they develop an algorithm using a semidefinite-programming (SDP) based rounding to produce a solution, which is then refined by the *local-search* procedure of (Tsourakakis et al., 2013). Their algorithm provides a 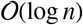 approximation guarantee, although in practice the approximation is shown to be much better. This is the algorithm we use in the extraction of the contrast subgraph.

### Appendix A.3. Computation of final solutions

The steps of our pipeline which lead to the final solutions are summarized in Figure 2C and Figure 2D. We next provide the technical details of these steps.

#### Bootstrap

Despite the classic setup of contrast subgraph problem aims to detect the one maximizing the contrast function, it is possible that 2 graphs can share more than a single contrastive structure. For this reason, we implement a bootstrap scheme in order to extract multiple solutions: this also helps us in preventing the possible bias brought by the imbalance in the groups. Therefore, we create 100 replications of the original dataset, each one containing, in equal number, the less represented class and a sample of the most-populated class. Computing the contrast subgraph in each of these datasets, we are able to extract more information. The multiple structures that we extract are finallky validated and reconciliated by means of a frequent itemset mining algorithm, that will be discussed later.

#### Choice of α

We next discuss the choice of the parameter α, necessary to compute the contrast subgraph according to the algorithm of (Lanciano et al., 2020). In general, the effective tuning of a parameter is a crucial aspect that comes out when designing a model. In order to address this, we follow an approach proposed by (De Vico Fallani et al., 2017) for the problem of thresholding, i.e., the determination of edges that are not significant or relevant, in a weighted graph.

(De Vico Fallani et al., 2017) introduces a density threshold able to filter out the weakest edges, by maximizing a quality function based on the ratio of the overall *efficiency*^3^ (Latora and Marchiori, 2001) of a network and its density. The quality function is the following:

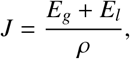

where *E_g_* represent the global efficiency, i.e. the average of the efficiency computed over all pair of nodes in the graph, *E_l_* is the local-efficiency of a network, i.e., the average of the global-efficiency computed over every induced ego-network by any node in the graph and *ρ* is the density of the network. By definition, these three quantities are normalized in the range [0, 1], and both the efficiency’s metrics are non-decreasing functions of *ρ*. This means that by the maximization of this function, we are able to output a solution that takes into account the natural trade-off generated by these quantities, and retain only those edges in the network that make it also either efficient and dense at the same time.

We find a strict connection between this framework and our necessity, since the parameter *α* is a penalty that assign a negative contribution to any edge inside the final solution of a contrast subgraph. Therefore, any edge (*u*, *v*) for which *w*(*u*, *v*) < *α* becomes a negative weighted edge that becomes detrimental for the objective function of the contrast subgraph algorithm, and we need to carefully choose the right value of *α* in order to not limit the choice of the algorithm for contrast subgraph, nor to make it shallow.

#### Statistical filtering

In the last step of the pipeline, we filter and reconciliate the multiple contrast subgraphs obtained due to the bootstrapping approach, using a filter based on the *maximal frequent itemset paradigm*. In this classic data mining task Luna et al. (2019), we are given a set of items *I* = {*i*_1_,…, *i_n_*}, a set of transactions *T* = {*t*_1_,…, *t_m_*} (*t_i_* ⊆ *I* ∀*i* ∈ {1,…*m*}) and scalars *k* and *s*, and we are required to find all the itemsets *i* ⊆ *I* s.t. |*i*| = *k* such that *s*(*i*) > *s*, where *s*(*i*) is the support of the item *i*, i.e. the number of transactions in which *i* is included.

In our setting, the given set of transactions are the contrast subgraphs, each one containing a set of different items (nodes), and we want to detect those items who are frequently co-occurring (those groups of ROIs that are often contrastive between the 2 different classes). Therefore, considering the different solutions computed, we mine the most frequent contrast subgraphs that are also statistically significant (such that False Discovery Rate ≤ 0.01), exploiting the proposal of (Kirsch et al., 2012), and retain only those that are maximals (i.e., sets that are not the subset of any other set in the collection).

More in details, (Kirsch et al., 2012) proposes an algorithm to perform a frequent itemset mining task with specific theoretical guarantees on the statistical significance of the output. This algorithm extracts frequent *k*-itemsets that are also statistically significant with False Discovery Rate (FDR) upper bounded by an input parameter *β*. This result relies on their proof that the number of k-itemsets with minimum support *s_min_* can be described with a Poisson distribution. Their algorithm start with the setting of *s_min_* according to a Monte-Carlo simulation that estimate a value for which for any support value above *s_min_* their approximation result hold. Obtained the frequent *k*-itemsets with support *s_min_* with any state-of-the-art algorithms, the algorithm tests for any frequent itemset the hypothesis that its support follows a Binomial distribution *Bin*(*n, p*), where *n* is equal to the number of transactions, and *p* is the empirical probability that the items appear together in a single transaction. Finally, computed the p-value of all the tests, the algorithm applies the multi-comparison test introduced by Benjamini and Yekutieli (Benjamini and Yekutieli, 2001) that bound the FDR given by the acceptance of the null hypothesis of multiple tests performed to a given parameter *β*.

In our pipeline, we apply this algorithm by considering the contrast subgraphs obtained in the boostrap procedure as the transactions, ranging *k* in the interval [2, *K*], where *K* represents the maximum size of a single transaction in the list, retaining only the connections statistically significant with a FDR less than 0.01.

### Appendix A.4. Coarse-graining of subgraphs and statistical selection of reduced edges

Consider a graph *G* defined on the node set of regions *V* and with edges *E_G_* ⊂ *V* × *V*, and a set of lobes *L* = {*L_i_*}_*i*_, such that each lobe, *L_i_* ∈ *L* is a subset of *V*, 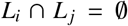, and ∪*L* = ∪*_i_L_i_* = *V*. We coarse-grain individual graphs to lobe-level graphs by defining a new lob graph *LG* with node set *L*, and edges *E_LG_*. Each edge *e* = (*u*, *v*) ∈ *E_G_* induces an edge in *e_lg_* = (*L_i_, L_j_*) ∈ *E_LG_* such that *u* ∈ *L_i_*, *v* ∈ *L_j_*. The weight of such edge *ω_e_l_g___* is the sum of the weight of the edges *e* that link the corresponding lobes *L_i_*, *L_j_*, *ω_e_l_g___* = (*L_i_*,*L_j_*) = ∑_*e*_=(*u,v*)∈*E_G_*|*u*∈*L_i_*,*v*∈*L_j_ ω_e_*. Note that it is possible for *u* and *v* to belong to the same lobe *L_i_* = *L_j_*, and therefore the graph *LG* can have self-loops with arbitrary weights. In fact, in the main text we use estensively the node selfloop weight (named node self-strength or integration).

For the hypo- and hyper-expressed results reported in Figure 4a, we perform an additional step before coarse-graining from regions to lobes. In particular, for each link *e* in the constrast subgraph for ASD(TD) subjects we compute its average weight 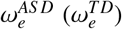 in the two groups, we build the distribution over the edges of the differences 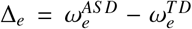, and we keep only the edges in with the top 10% largest difference. We consider these edges hyper-expressed, and then coarse-grained the resulting graph to lobe level. For the hypo-expressed we perform the same procedure keeping the edges corresponding to the smallest 10% of differences.

## Footnotes

1 http://fcon_1000.projects.nitrc.org/indi/abide/

2 http://www.preprocessed-connectomes-project.org/abide

3 The efficiency of a pair of nodes in a graph is computed as the multiplicative inverse of the shortest path distance between the nodes.

## Notes

### Competing Interest Statement

The authors have declared no competing interest.

